# Principal Component Analysis of Major Biometric Traits and Indices of Indigenous Sheep Ewe Populations, and Relationships with their Body Weights

**DOI:** 10.1101/2025.01.28.635228

**Authors:** Teweldemedhn Mekonnen, Medhanye Araya, Solomon Tesfahun

## Abstract

The field data collection was performed before the war (before October 2020) in Tigray Region, Ethiopia. The principal component analysis (PCA) is a grouping of variables or dimension reduction technique. The objective of this study was to characterize and identify the most underlying biometric traits and biometric indices of the indigenous sheep ewe populations, and determine the relationships of morphometric traits and morphometric indices with the body weights of the ewe populations. Sample animals of Begait (160), Rutanna (129) and Arado (156) ewe populations which totaled 445 with permanent pair of incisors (1PPI-4PPI) were randomly involved in the field data collection. Statistical Package for Social Sciences software was used for statistical data analysis. The PCA of the morphometric traits (09) extracted two principal components (PCs) in Begait ewes, two PCs in Rutanna ewes and one PC in Arado ewes, and the PCA of the morphometric indices (20) extracted five PCs in Begait ewes, six PCs in Rutanna ewes and five PCs in Arado ewes. The compact index mainly indicated that Rutanna and Begait ewes are larger in size and suitable for mutton production whilst Arado ewes are suitable for milk production. The correlations among the biometric traits and the body weights of the ewe populations were highly significantly (*P*<0.01) different except the correlation between body weight and rump length of Rutanna ewes (*P*<0.05). The biometric indices which consisted of height index, cephalic index, longitudinal pelvic index, height slope, over increase index, body ratio, foreleg length of the three ewe populations and their body weights were with little or no correlations (*P*>0.05). Genetic characterizations of the indigenous (Begait and Arado) and the transboundary (Rutanna) sheep populations should be complemented to confirm and distinguish the purpose of the populations.

## INTRODUCTION

Ethiopia is a center of livestock diversity because the country is a route for migration from Asia into Africa (Solomon, 2008). Rutanna sheep is a desert sheep in the Sudan, and are kept mainly for meat production. Rutanna breed of sheep is preferred in border markets in Western Ethiopia for export due to its higher growth potential and large body size (Ali, 2003; Mohammed, 2015). Rutanna sheep is one of the thin-tailed sheep which is mainly found in Sudan, and the sheep is also distributed in the border areas of Ethiopia.

Extensive management systems could be the causes of the loss of valuable genes due to the practices of uncontrolled breeding (Groeneveld *et al*., 2010). The phenotypic characterization of the observable characteristics of livestock breed(s) is used to identify and document within and between breed variation. Livestock planning on improvement, sustainable utilization, conservation strategies and breeding programs depend partly on their morphometric characterizations (FAO, 2012). Body measurements and body indices are used for breed characterizations in goats and sheep (Marković *et al*., 2019; Putra and Ilham, 2019). Larger differences of body measurements are indicators of larger distances among breeds (Markovic *et al*., 2019). Body size data collection is relatively easier to get and cheap when compared to protein or DNA data collection (Handiwirawan *et al*., 2011). Linear body (biometric) measurements can be used as an option of selection criteria for improvement of meat production (Verma *et al*., 2015; Putra and Ilham, 2019). Many researchers have been traditionally used morphological measurements for characterization of native sheep breeds. The relationship between body conformation and function of different animal species and breeds has been widely observed (Latorre *et al*., 2011). Body size and shape are important traits in meat animals, and the characterization of local genetic resources depends on the knowledge of the variation of morphological traits which are fundamental in classification of livestock breeds based on size and shape (Ferra *et al*., 2010; Agga *et al*., 2010; Leng *et al*., 2010; Yakubu, 2010a and 2010b).

Multivariate statistical tools or methods such as principal component analysis (PCA) is a more reliable assessment of morphometric relationship among livestock breeds (Yakubu, 2013). PCA is a multivariate technique of statistical method which could be used with success when morphological variables are interdependent, and principal components (PCs) are a weighted linear combination of correlated variables, explaining a maximal amount of variance of the variables (Truxillo, 2003). The PCA aids in data reduction and breaks multicollinearity which may lead to wrong inferences (Yakubu and Ayoade, 2009). PCA is an interdependence technique whose primary purpose is to define the underlying structure among the variables under study. First few components account for the highest proportion of the total variance though the number of components generated in PCA equals the number of variables in the study. The PCA has been used as a tool in the assessment of the body conformation which can be conducted to understand the complex growth process in the body dimensions of an animal during growth period. PCA results not only impact the management of animals but also help in conservation and selection of multiple traits by breeders (Salako, 2006; Khargharia *et al*., 2015). PCA analysis was used to extract factors contributing to variation among individual animals based on their body measurements (Salako, 2006; Mavule *et al*., 2013; Yakubu, 2013; Yunusa *et al*., 2013; Khargharia *et al*., 2015).

The aesthetics and production potential of the animals are assessed by the body structural indices (Banerjee, 2017). The index system was first developed for the assessment of the type and function of cattle populations and the system was suggested to other species (Alderson, 1999). Structural indices or morphological indices are used to assess the type and function of the sheep population (Maciejowski and Zieba, 1982; Chacon *et al*., 2011). The morphological or structural indices give information about breed characteristics in terms of structure and proportions on functional traits of animals ((Esquivelzeta *et al*., 2011). Individual measurement traits are less accurate estimation of an animal’s conformation than morphological indices. Structural or morphometric indices provide tested empirical values (no single measurements) and used for the assessment of type, weight and function, and boost the ability of breeders to select potential breeding stock (Salako, 2006). Morphometric measurements are related with production characteristics. Structural indices are superior over single measurements and provide evaluation of animals to buyers (Mohammed and Amin, 1997). Morphometric indices are calculated from the morphometric traits and are generally used for the estimation of conformation of animals (Mohammed and Amin, 1997; Pares- Casanova *et al*., 2013; Dauda, 2018).

There are indigenous sheep populations (Begait and Arado) and a transboundary sheep (Rutanna) in Western Zone of Tigray, Ethiopia. These three sheep populations contribute in the social, economic and cultural issues of the communities in the Zone. However, the communities lack knowledge on biometric indices for the proper identification of the type and function of the sheep populations due to lack of information on the biometric indices of the indigenous sheep of the study area. Biometric indices greatly help the commercialized small ruminant breeders in identifying whether a breed is used for meat/mutton production or milk production or dual purpose breed. The biometric indices are mainly used in developing animal breeding strategies and animal selection (Diriba, 2020). Hence, this study will also be used for the research and development institutions. The biometric indices are simple mathematical calculations from the morphometric traits and body weight, and used for individuals involved in small ruminant fattening and breeding. Therefore, the objective of this study was to characterize and identify the most underlying morphometric traits and morphometric indices of the indigenous ewe populations, and determine the relationships of morphometric traits and morphometric indices with the body weights of the ewe populations.

## MATERIALS AND METHODS

### Description of the Study Areas

Kafta Humera, Tsegede and Welkait districts were the field data collection areas where the indigenous sheep (Begait and Arado ewes) and transboundary sheep (Rutanna ewes) are available. Kafta Humera district is the lowland part of Western Zone of Tigray Region, Ethiopia whereas Welkait and Tsegede districts are the highland areas of Western Zone of Tigray Regional State. The Western Zone of Tigray is located at 570 and 991 kilometers (Km) far from Mekelle and Addis Ababa, respectively (ZOIC, 2015). The Zone also lies at 13°42′ to 14°28′ North latitudes and 36°23′ to 37°31′ East longitudes (Mekonnen *et al*., 2011). The altitude, rainfall, temperature and non-arable land uses of Kafta Humera, Welkait and Tsegede districts are presented (Table 1).

**Table 1.** Agro-climatic and non-arable land use of the study districts.

| **Agro-climate and land use** | **Kafta Humera** | **Welkait** | **Tsegede** |
| --- | --- | --- | --- |
| Altitude (MASL) | 500-1849 | 700-2354 | 680-3008 |
| **Agro-ecology (%)** |  |  |  |
| Lowland (*Kola*) | 86 | 60 | 70 |
| Midland (*Weina dega*) | 14 | 40 | 22 |
| Highland (*Dega*) | - | - | 9 |
| Rainfall (mm) | 650-750 | 700-1800 | 1200-2500 |
| Temperature (°C) | 25-48 | 18-25 | 12-35 |
| **Non-arable land uses (%)** |  |  |  |
| Forestry land | 33 | 19 | 35 |
| Pastureland | 5 | 18 | 22 |

Meter Above Sea Level (MASL), millimeter (mm), Source: Tesfay *et al.*, 2019

### Data collection and statistical analysis

Begait (160), Rutanna (129) and Arado (156) adult sheep which totaled 445 sample animals were randomly involved in the field data collection. The ages of the animals were determined by dentition and were aged from one permanent incisor (1PPI) up to four permanent incisors (4PPI) (Figure 1).

**Figure 1.**
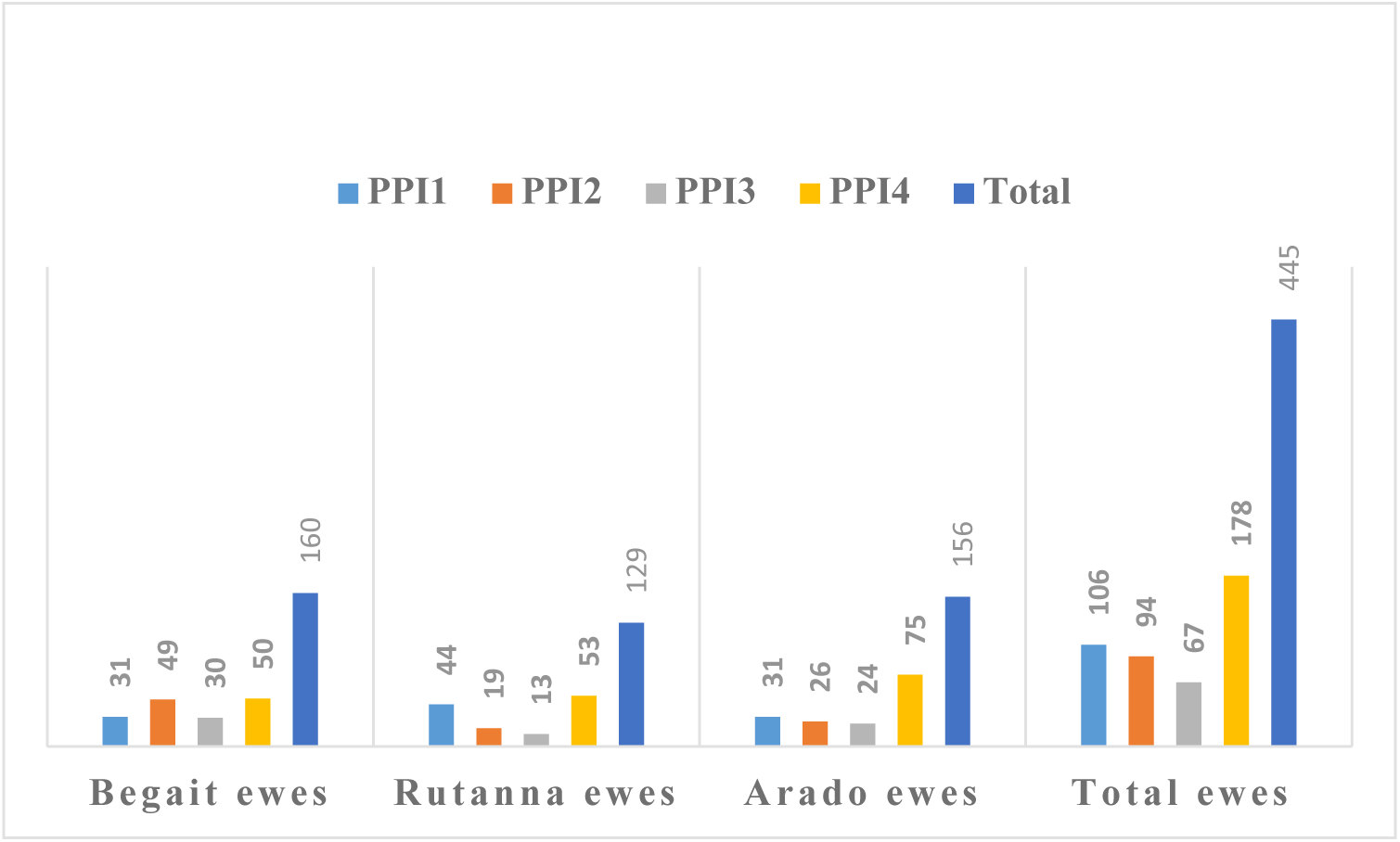
Number of observations of ewes by breed and age class used.

The indigenous sheep populations were kept at low input extensive production system. The quantitative data were collected according to the guideline of FAO (2012) standard breed descriptor list of sheep breeds. The linear body traits were measured using flexible measuring tape while body weight (BW) was measured using suspended spring balance having 100 Kg capacity.

Nine (09) linear body (morphometric) traits were measured (in centimeter) which comprised of Body length (BL), Heart Girth or Chest circumference or Chest girth (ChG), Chest depth (ChD), Height at withers (HW), Rump height (RuH), Rump length (RuL), Rump width (RuW), Head length (HdL), and Head width (HdW), therefore, there were ten biometric traits data collected. Twenty (20) morphometric indices were analyzed which comprised of Length index (LI), height index (HI), Depth index (DI), Relative depth of thorax (RDT), Thoracic development index (TDI), conformation index (CI), Compact index (CoI), Body weight index (BWI), Body index (BI), Proportionality index (PrI), Area index (AI), Cephalic index (CpI), Pelvic index (PI), Transverse pelvic index (TPI), Longitudinal pelvic index (LPI), Height slope (HS), Over increase index (OII), Body ratio (BR), Rump length index (RLI), and Foreleg length (FL).

Principal Component Analysis or PCA (FAO, 2012) was used as a statistical method of data analysis. **Correlation matrix** of PCA of the ewe populations was used to identify the principal components in the morphometric traits and indices of the ewe populations. PCA data set validity test was computed using Kaiser-Meyer-Olkin (KMO) measure of sampling adequacy and Bartlett’s test of sphericity. The KMO measure of sampling adequacy vary between zero and one, and the value close to zero indicates that there are *large partial correlations compared to the total correlations* whilst the value close to one indicates that there is appropriate and adequate sampling. Factor analysis is conducted when the KMO measure of sampling adequacy is greater than 0.50 (Putra and Ilham, 2019), and the KMOs of the morphometric traits and morphometric indices of this study are presented (Table 2). The proportion of variance of the variable which jointly measure and explained by all other factors is the estimate of communality for each variable (Verma *et al*., 2015). Statistical Package for Social Sciences (SPSS, 2017) software was used for statistical data analysis. The biometric traits (10) and the biometric indices (20) were summarized by descriptive statistics. One-way ANOVA (Tukey at Alpha=0.05) was used to compare the biometric indices among the ewe populations.

**Table 2.** KMO and Bartlett’s Test of Sphericity.

| Measure | Begait ewes |  | Rutanna ewes |  | Arado ewes |  |
| --- | --- | --- | --- | --- | --- | --- |
|  | Morphometric traits | Morphometric indices | Morphometric traits | Morphometric indices | Morphometric traits | Morphometric indices |
| KMO | 0.85 | 0.71 | 0.82 | 0.69 | 0.86 | 0.68 |
| Bartlett's test of sphericity | 0.000 | 0.000 | 0.000 | 0.000 | 0.000 | 0.000 |
| (P-value) |  |  |  |  |  |  |

The biometric or morphometric or body structural indices (Table 3) were calculated according to the formulas of different authors to describe the type and function of the indigenous ewes in the study area.

**Table 3.** Formulas for the calculation of the morphometric/body structural indices.

| Body structural indices | Formula | Description | Source(s) |
| --- | --- | --- | --- |
| Length index (LI) or relative body index (RBI) | $LI = \frac{BL}{HW}$ | | Salako, 2006; Handiwirawan *et al.*, 2011; Dauda, 2018a; Dauda, 2018b; Putra and Ilham, 2019; Birara *et al.*, 2021; Ilham *et al.*, 2023 |
| Body proportion index (BPI) or height index (HI) | $HI = \frac{HW}{RuH} \times 100$ | | Ibrahim *et al.*, 2023 |
| Depth index (DI) | $DI = \frac{ChD}{HW}$ | | Salako, 2006; Handiwirawan *et al.*, 2011; Putra and Ilham, 2019; Ilham *et al.*, 2023 |
| Relative depth of thorax (RDT) | $RDT = \frac{ChD}{HW} \times 100$ | | Chiemela *et al.*, 2016; Putra and Ilham, 2019; Birara *et al.*, 2021; Ibrahim *et al.*, 2023; Ilham *et al.*, 2023 |
| Thoracic development index (TDI) | $TDI = \frac{ChG}{HW}$ | TDI with values above 1.2 indicating animal with good TD | Chiemela *et al.*, 2016; Dauda, 2018a; Dauda, 2018b; Putra and Ilham, 2019; Birara *et al.*, 2021; Ilham *et al.*, 2023 |
| Baron and Crevat index (BCI) or conformation index (CI) | $CI = \left(\frac{ChG}{HW}\right) \times 2$ | CI should be close to 2.1. The bigger the CI, the closer the animal is to the traction type; the smaller the CI, the weaker the animal will be. | Updated |
| Compact index (CoI) | $CoI = \left(\frac{BW}{ChD}\right) \times 3$ | Compact index indicates how compact the animal is. Meat type animals have values above 3.15. Value close to 2.75 indicates dual purpose and close to 2.60 indicates that the animals are more suitable for milk purpose. | Updated |
| Body weight index (BWI) | $BWI = \frac{BW}{HW} \times 100$ | | Boujenane, 2015; Khargharia *et al.*, 2015; Dauda, 2018b; Marković *et al.*, 2019; Depison *et al.*, 2020 |
| Body index (BI) | $BI = \frac{BL}{ChG}$ | When BI is greater than 0.90, the animal is ***longiline***; between 0.86 to 0.88 is ***medigline***; and less than 0.85, it is ***brevigline*** | Updated |
| Proportionality index (PrI) | $PrI = \frac{HW}{BL} \times 100$ | | Chiemela *et al.*, 2016; Dauda, 2018a; Dauda, 2018b; Putra and Ilham, 2019; Depison *et al.*, 2020; Birara *et al.*, 2021; Ibrahim *et al.*, 2023; Ilham *et al.*, 2023 |
| Area index (AI) | $AI = HW \times BL$ | | Dauda, 2018a; Dauda, 2018b; Putra and Ilham, 2019; Depison *et al.*, 2020; Birara *et al.*, 2021; Ilham *et al.*, 2023 |
| Cephalic index (CpI) | $CpI = \frac{HdW}{HdL} \times 100$ | | Chiemela *et al.*, 2016; Putra and Ilham, 2019 |
| Pelvic index (PI) | $PI = \frac{RuW}{RuL} \times 100$ | | Chiemela *et al.*, 2016; Dauda, 2018a; Dauda, 2018b; Birara *et al.*, 2021 |
| Transverse pelvic index (TPI) | $TPI = \frac{RuW}{RuH} \times 100$ | | Chiemela *et al.*, 2016; Birara *et al.*, 2021 |
| Longitudinal pelvic index (LPI) | $LPI = \frac{RuL}{RuH} \times 100$ | | Chiemela *et al.*, 2016; Birara *et al.*, 2021 |
| Height slope (HS) | $HS = HW - RuH$ | | Salako, 2006; Handiwirawan *et al.*, 2011 |
| Over increase index (OII) | $OII = \frac{RuH}{HW} \times 100$ | | Cited in Diriba (2020) |
| Body ratio (BR) | $BR = \frac{HW}{RuH}$ | | Chiemela *et al.*, 2016; Birara *et al.*, 2021; Ilham *et al.*, 2023 |
| Rump length index (RLI) | $RLI = \frac{RuL}{BL} \times 100$ | | Cited in Diriba (2020) |
| Foreleg length (FL) | $FL = HW - ChD$ | | Salako, 2006; Handiwirawan *et al.*, 2011; Ilham *et al.*, 2023 |

### Ethical Statement Approval

There is no local or national animal research ethics committee of live animal external body parts data measurement and use. Hence, there was no ethical approval used because the external animal body parts were measured without animal slaughtering or any injury and made under the consent of their owners. The morphometric traits of the animal body parts were measured early in the morning (FAO, 2012) and were measured humanely.

## RESULTS AND DISCUSSION

### Morphometric traits of indigenous sheep ewe populations

The biometric traits comprise of the morphometric (linear body) traits and body weights of the ewe populations. The major morphometric traits were used in phenotyping the indigenous sheep ewe populations. The mean (cm) body length (BL), chest girth (ChG), rump height (RuH) and body weight (BW) in Kg of Begait (72.38, 76.52, 75.16 and 39.46), Rutanna (72.98, 80.42, 77.29 and 45.27) and Arado (66.98, 69.94, 71.09 and 28.04) ewe populations were not comparable, respectively (Table 4). These three indigenous sheep exhibited higher RuH than HW which is in line with Salako (2006) report on West African Dwarf (WAD) and Yankasa sheep of Nigeria. The biometric traits revealed that Arado ewes are smaller in size than the size of Begait and Rutanna ewes because Arado ewes exhibited shorter BL (66.48), HW (69.99) and RuH (71.09 cm), and smaller body size (28.04 Kg). The basis of the variation could be due to mainly their variation in genetic makeup. The BW of Begait, Rutanna and Arado ewes are not similar with the BW of WAD (25.03 Kg) and Yankasa (41.60 Kg) sheep in Nigeria (Salako, 2006). The difference might be due to genotype, environmental and the interaction effects of genotype-environment.

**Table 4.** Major morphometric (biometric) traits of indigenous sheep ewe (n =445) populations.

| **LBT** | **Begait ewes (n=160)** |  |  | **Rutanna ewes (n=129)** |  |  | **Arado ewes (n=156)** |  |  |
| --- | --- | --- | --- | --- | --- | --- | --- | --- | --- |
|  | **Mean** | **SD** | **Variance** | **Mean** | **SD** | **Variance** | **Mean** | **SD** | **Variance** |
| BL | 72.38 | 3.07 | 9.41 | 72.98 | 3.69 | 13.69 | 66.48 | 2.88 | 8.32 |
| ChG | 76.52 | 3.84 | 14.71 | 80.42 | 5.09 | 25.86 | 69.94 | 3.57 | 12.74 |
| ChD | 37.34 | 1.89 | 3.59 | 38.25 | 2.20 | 4.84 | 34.43 | 1.68 | 2.83 |
| HW | 74.37 | 3.04 | 9.25 | 76.47 | 3.25 | 10.58 | 69.99 | 2.73 | 7.44 |
| RuH | 75.16 | 3.06 | 9.38 | 77.29 | 3.25 | 10.58 | 71.09 | 2.76 | 7.59 |
| RuL | 18.69 | 1.34 | 1.79 | 18.57 | 1.40 | 1.97 | 14.84 | 1.15 | 1.32 |
| RuW | 17.32 | 1.02 | 1.04 | 17.33 | 1.52 | 2.29 | 16.10 | 0.97 | 0.93 |
| HdL | 23.03 | 0.94 | 0.89 | 22.61 | 1.07 | 1.15 | 21.76 | 0.84 | 0.70 |
| HdW | 13.36 | 0.73 | 0.53 | 12.59 | 0.78 | 0.60 | 11.69 | 0.51 | 0.26 |
| BW | 39.46 | 5.57 | 31.06 | 45.27 | 8.00 | 64.06 | 28.04 | 3.45 | 11.87 |

Numbers of ewes (n), Linear Body Traits (LBT), Standard Deviation (SD), Body length (BL), Chest girth (ChG), Chest depth (ChD), Height at withers (HW), Rump height (RuH), Rump length (RuL), Rump width (RuW), Head length (HdL), Head width (HdW), Body weight (BW)

The present mean BL, HW and RuH of Arado ewes are in line with the BL (67.30), HW (70.30) and RuH (71.42) of Nigerian sheep (Olaniyi *et al*., 2018), however, the BW (25.07) of Nigerian sheep is less than the BW of Arado ewes whilst the ChG (79.89) of Nigerian sheep is larger than the present ChG (69.94) of Arado ewes. The differences in BW and ChG could be due to genotype, environmental and their interaction effects. Moreover, except ChG, the HW, RuH and BL of Arado ewes are not similar with the HW (61.74), RuH (59.20) and BL (62.56) of WAD sheep (Salako, 2006). The BL and HW of Begait and Rutanna ewes are similar with the BL and HW of Yankasa sheep in Nigeria (Salako, 2006), however, the ChG and RuH of Begait and Rutanna ewes are not similar with the ChG (86.63) and RuH (72.57) of Yankasa sheep (Salako, 2006). The present mean BL of Begait and Rutanna ewes are similar with the BL of Wonosobo (71.14) sheep and Batur (72.27) sheep (Ibrahim *et al*., 2023) whilst the HW and RuH of Begait, Rutanna and Arado ewes are not similar with the HW and RuH of Wonosobo (64.79, 63.79) sheep and Batur (63.44, 63.71) sheep (Ibrahim *et al*., 2023), respectively. All the differences could be due to genotype, environmental and their interaction effects. The RuH of Begait, Rutanna and Arado ewes are not similar with the RuH of Corriedale (65.92) and Australian Merino (68.45) sheep (Latorre *et al*., 2011) due to their differences in genetic makeup, environment and the interaction effect of the genotype and the environment. However, there are similarities of the BL of Begait and Rutanna ewes and the BL of Australian Merino (74.85), BL of Arado ewes and the BL of the Corriedale (65.90) sheep, and the HW of Arado ewes and the HW of the Australian Merino (67.25).

The correlation coefficients of the morphometric traits and body weights of the indigenous sheep ewe populations were highly significantly (*P*<0.01) different (Table 5). The strong correlations indicate that the selection of the morphometric traits could improve the body weight of the ewes. There were little or no correlation among some morphometric traits which included RuW*HdW (0.100 in Begait), HdL*RuL (0.083 in Rutanna) and RuL*ChG (0.166 in Rutanna) of the indigenous sheep ewe populations (Table 5). Except the correlation coefficient of BW*RuL (*P*<0.05) in Rutanna ewes, the correlation coefficients of all morphometric traits and body weights of the three ewe populations were highly significantly (*P*<0.01) different (Table 5). The communalities of morphometric traits and morphometric indices of the indigenous ewe populations are also presented (Table 6).

**Table 5.** Pearson’s correlation coefficient of the morphometric traits and body weights of the indigenous sheep ewe populations (Top matrix- Begait, Middle matrix- Rutanna and Bottom matrix- Arado sheep)

| LBT | BL | ChG | ChD | HW | RuH | RuL | RuW | HdL | HdW | BW |
| --- | --- | --- | --- | --- | --- | --- | --- | --- | --- | --- |
| BL | 1 | 0.555** | 0.683** | 0.575** | 0.525** | 0.238** | 0.402** | 0.495** | 0.265** | 0.590** |
| ChG | 0.597** | 1 | 0.831** | 0.533** | 0.519** | 0.325** | 0.523** | 0.482** | 0.370** | 0.850** |
| ChD | 0.551** | 0.725** | 1 | 0.614** | 0.596** | 0.323** | 0.497** | 0.522** | 0.343** | 0.744** |
| HW | 0.468** | 0.543** | 0.510** | 1 | 0.809** | 0.315** | 0.406** | 0.534** | 0.406** | 0.501** |
| RuH | 0.396** | 0.505** | 0.495** | 0.789** | 1 | 0.290** | 0.357** | 0.548** | 0.498** | 0.502** |
| RuL | 0.182* | 0.166 | 0.281** | 0.250** | 0.334** | 1 | 0.272** | 0.213** | 0.218** | 0.216** |
| RuW | 0.325** | 0.239** | 0.463** | 0.270** | 0.287** | 0.398** | 1 | 0.292** | 0.100 | 0.507** |
| HdL | 0.418** | 0.620** | 0.562** | 0.452** | 0.445** | 0.083 | 0.267** | 1 | 0.478** | 0.465** |
| HdW | 0.338** | 0.371** | 0.435** | 0.280** | 0.229** | 0.316** | 0.353** | 0.442** | 1 | 0.307** |
| BW | 0.659** | 0.872** | 0.708** | 0.458** | 0.453** | 0.192* | 0.322** | 0.606** | 0.401** | 1 |
| **BL** | 1 |  |  |  |  |  |  |  |  |  |
| **ChG** | 0.591** | 1 |  |  |  |  |  |  |  |  |
| **ChD** | 0.586** | 0.739** | 1 |  |  |  |  |  |  |  |
| **HW** | 0.443** | 0.580** | 0.584** | 1 |  |  |  |  |  |  |
| **RuH** | 0.426** | 0.483** | 0.544** | 0.807** | 1 |  |  |  |  |  |
| **RuL** | 0.337** | 0.369** | 0.336** | 0.356** | 0.294** | 1 |  |  |  |  |
| **RuW** | 0.506** | 0.488** | 0.509** | 0.314** | 0.263** | 0.189* | 1 |  |  |  |
| **HdL** | 0.463** | 0.590** | 0.487** | 0.494** | 0.381** | 0.468** | 0.414** | 1 |  |  |
| **HdW** | 0.436** | 0.430** | 0.418** | 0.443** | 0.358** | 0.212** | 0.344** | 0.428** | 1 |  |
| **BW** | 0.590** | 0.761** | 0.638** | 0.447** | 0.419** | 0.213** | 0.524** | 0.502** | 0.396** | 1 |

\*\* . Correlation is significant at the 0.01 level (2-tailed). \* . Correlation is significant at the 0.05 level (2-tailed).

**Table 6.**
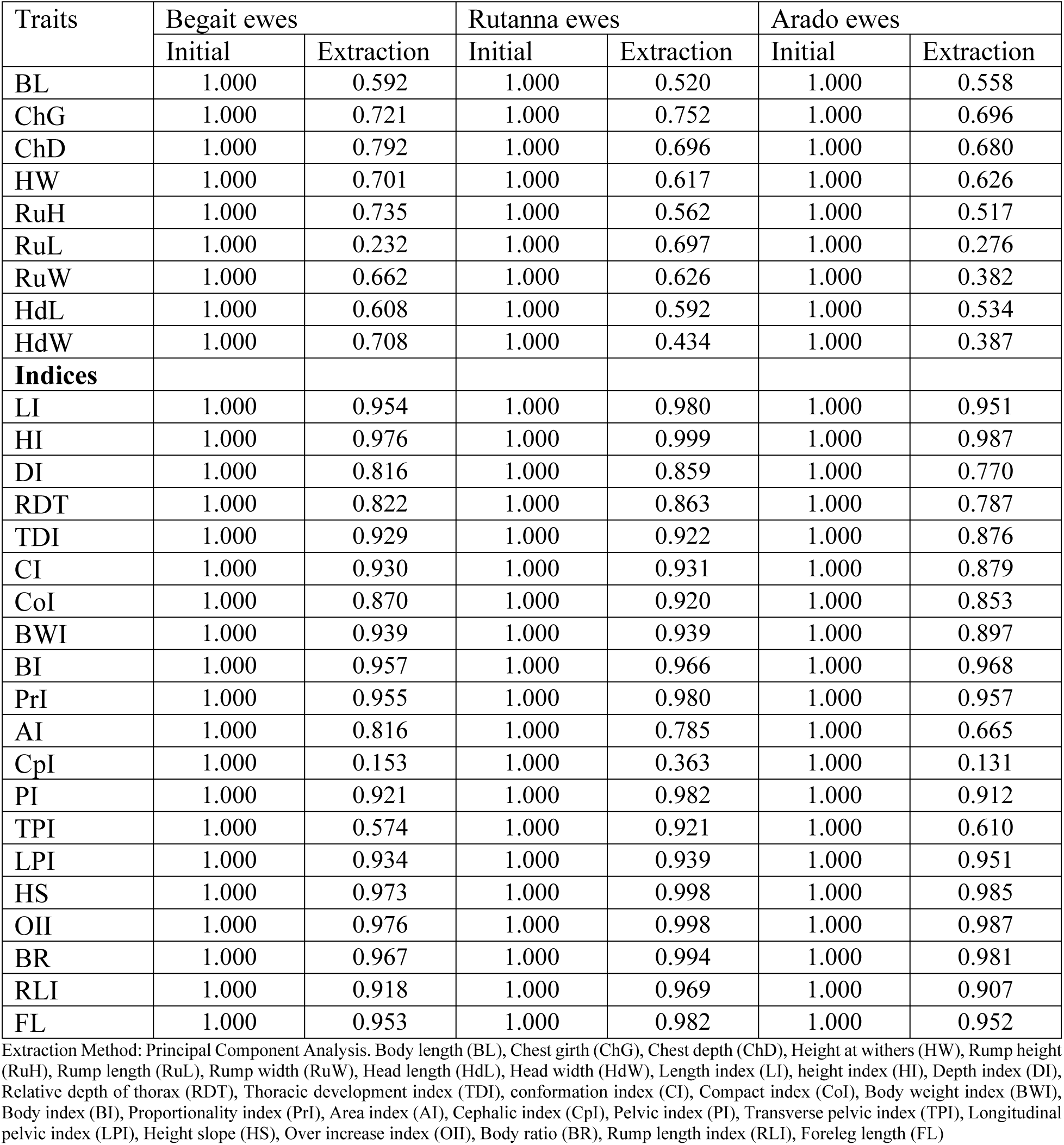
Communalities (morphometric traits and morphometric indices) of ewe populations.

Total variance explained (Table 7) and rotated component matrix (Table 8) of the morphometric traits of the ewe populations are clearly reported. Morphometric traits with total initial eigenvalues greater than one were extracted as principal components (PCs). Hence, there were two PCs extracted which accounted for 63.9% of the total variance (Begait ewes), two PCs extracted which accounted for 61.1% of the total variance (Rutanna ewes) and one PC extracted which accounted for 51.7% of the total variance (Arado ewes); the PCs were extracted from the different morphometric traits of the ewe populations (Tables 7 and 8). Components with eigenvalues greater than one are rotated using the varimax rotation. However, the rotation sums of squared loadings of morphometric traits of Arado ewes with eigenvalue of greater than one (4.656) was absent because there was only one PC extracted from the morphometric traits of Arado ewes and the solution could not be rotated. Not the rotated component matrix of Arado ewes is reported but component matrix of the principal component (PC) of the morphometric traits of Arado ewes is presented (Table 8).

**Table 7.**
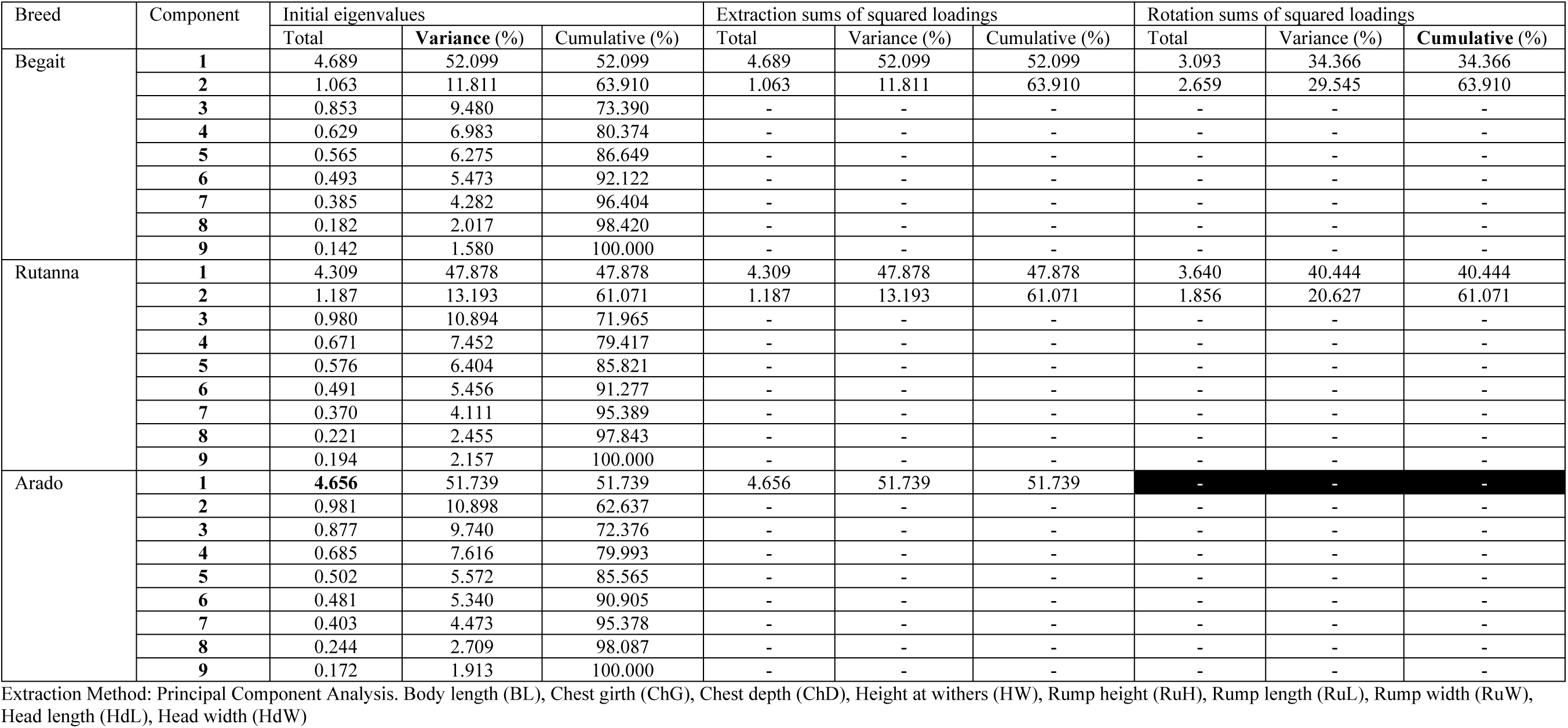
Total variance explained (biometric or morphometric traits)

| Breed | Component | Initial eigenvalues |  |  | Extraction sums of squared loadings |  |  | Rotation sums of squared loadings |  |  |
| --- | --- | --- | --- | --- | --- | --- | --- | --- | --- | --- |
|  |  | Total | Variance (%) | Cumulative (%) | Total | Variance (%) | Cumulative (%) | Total | Variance (%) | Cumulative (%) |
| Begait | 1 | 4.689 | 52.099 | 52.099 | 4.689 | 52.099 | 52.099 | 3.093 | 34.366 | 34.366 |
|  | 2 | 1.063 | 11.811 | 63.910 | 1.063 | 11.811 | 63.910 | 2.659 | 29.545 | 63.910 |
|  | 3 | 0.853 | 9.480 | 73.390 | - | - | - | - | - | - |
|  | 4 | 0.629 | 6.983 | 80.374 | - | - | - | - | - | - |
|  | 5 | 0.565 | 6.275 | 86.649 | - | - | - | - | - | - |
|  | 6 | 0.493 | 5.473 | 92.122 | - | - | - | - | - | - |
|  | 7 | 0.385 | 4.282 | 96.404 | - | - | - | - | - | - |
|  | 8 | 0.182 | 2.017 | 98.420 | - | - | - | - | - | - |
|  | 9 | 0.142 | 1.580 | 100.000 | - | - | - | - | - | - |
| Rutanna | 1 | 4.309 | 47.878 | 47.878 | 4.309 | 47.878 | 47.878 | 3.640 | 40.444 | 40.444 |
|  | 2 | 1.187 | 13.193 | 61.071 | 1.187 | 13.193 | 61.071 | 1.856 | 20.627 | 61.071 |
|  | 3 | 0.980 | 10.894 | 71.965 | - | - | - | - | - | - |
|  | 4 | 0.671 | 7.452 | 79.417 | - | - | - | - | - | - |
|  | 5 | 0.576 | 6.404 | 85.821 | - | - | - | - | - | - |
|  | 6 | 0.491 | 5.456 | 91.277 | - | - | - | - | - | - |
|  | 7 | 0.370 | 4.111 | 95.389 | - | - | - | - | - | - |
|  | 8 | 0.221 | 2.455 | 97.843 | - | - | - | - | - | - |
|  | 9 | 0.194 | 2.157 | 100.000 | - | - | - | - | - | - |
| Arado | 1 | **4.656** | 51.739 | 51.739 | 4.656 | 51.739 | 51.739 | - | - | - |
|  | 2 | 0.981 | 10.898 | 62.637 | - | - | - | - | - | - |
|  | 3 | 0.877 | 9.740 | 72.376 | - | - | - | - | - | - |
|  | 4 | 0.685 | 7.616 | 79.993 | - | - | - | - | - | - |
|  | 5 | 0.502 | 5.572 | 85.565 | - | - | - | - | - | - |
|  | 6 | 0.481 | 5.340 | 90.905 | - | - | - | - | - | - |
|  | 7 | 0.403 | 4.473 | 95.378 | - | - | - | - | - | - |
|  | 8 | 0.244 | 2.709 | 98.087 | - | - | - | - | - | - |
|  | 9 | 0.172 | 1.913 | 100.000 | - | - | - | - | - | - |

Extraction Method: Principal Component Analysis. Body length (BL), Chest girth (ChG), Chest depth (ChD), Height at withers (HW), Rump height (RuH), Rump length (RuL), Rump width (RuW), Head length (HdL), Head width (HdW)

**Table 8.**
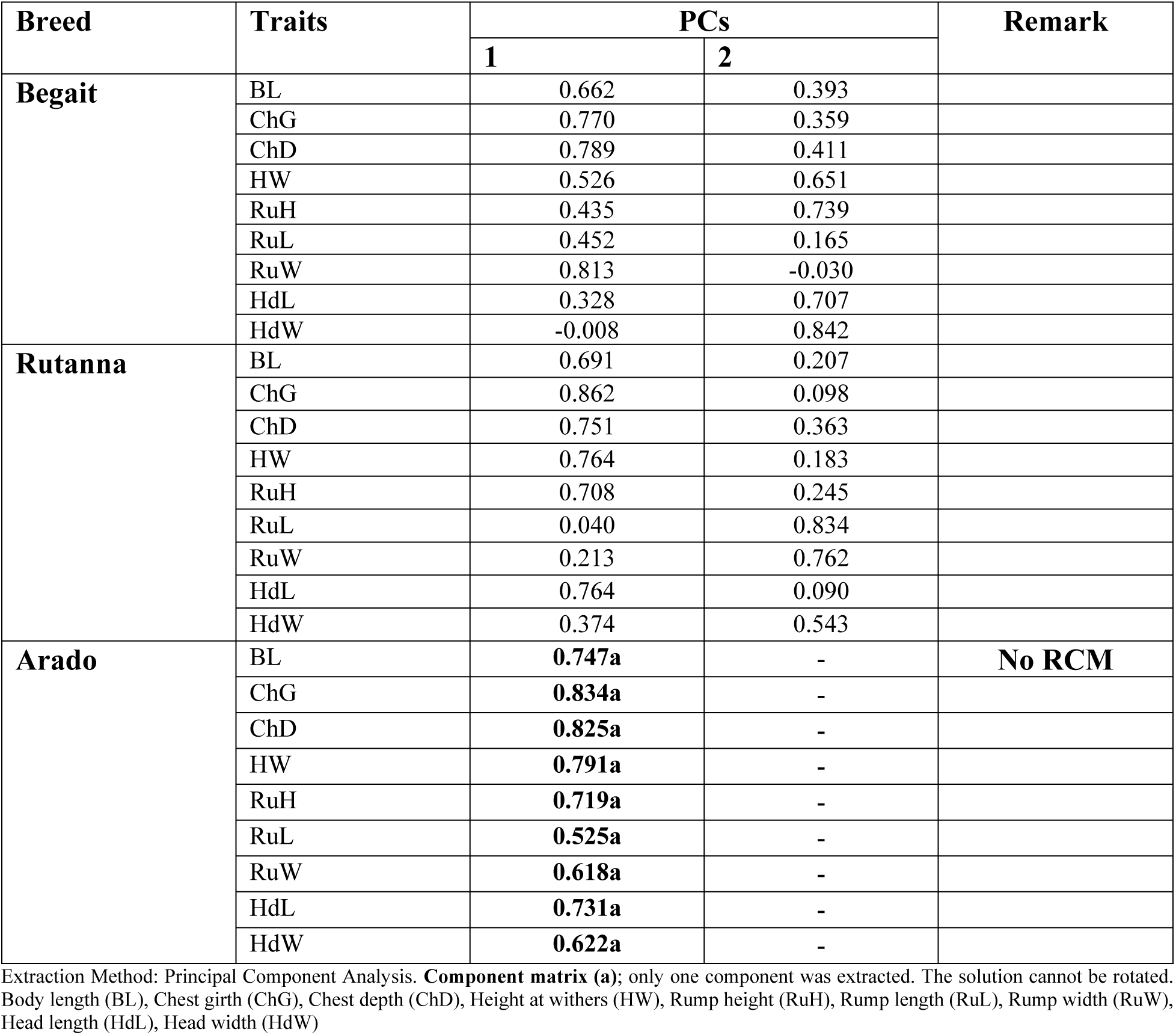
Rotated component matrix (RCM) of different principal components (PCs) of morphometric traits of different sheep ewe populations.

### Biometric (Morphometric or body structural) indices of indigenous sheep ewe populations

The morphometric indices or structural indices also helped in phenotyping the indigenous sheep ewe populations because except few, the biometric indices were significantly (*P*<0.05) different among breeds (Table 9). The means of Begait and Rutanna ewes of some morphometric (biometric) indices were length index-LI (0.97, 0.96), height index-HI (98.99, 98.97), conformation index-CI (2.10, 2.10), compact index-CoI (3.16, 3.54), body weight index-BWI (52.99, 59.11), body index-BI (0.95, 0.91), area index-AI (5388.5, 5586.4) and body ratio-BR (0.99, 0.99) whilst the mean of LI (0.95), HI (98.49), CI (1.99), CoI (2.44), BWI (40.03), BI (0.95), AI (4656.7) and BR (0.99) of Arado ewes were also displayed (Table 9). The BI of Rutanna ewes was slightly lower than the BI of the Begait and Arado ewes due to the influence of genetic and non-genetic factors (Chacon *et al*., 2011). The over increase index-OII (101.60), LI and BR of Arado ewes are in agreement with the OII (101.88), LI (0.96) and BR (0.98) of Nigerian sheep (Olaniyi *et al*., 2018) whereas the cephalic index-CpI (53.75), HI and BI of Arado ewes are not similar with the CpI (76.71), HI (106.15) and BI (84.81) of Nigerian sheep (Olaniyi *et al*., 2018). The differences might be due to genotype, environment, the interaction effect of genotype and environment, sample size and field measurement errors. The HI of Begait, Rutanna and Arado ewes are in agreement with the HI of Batur (99.51) sheep in Indonesia (Ibrahim *et al*., 2023).

**Table 9.**
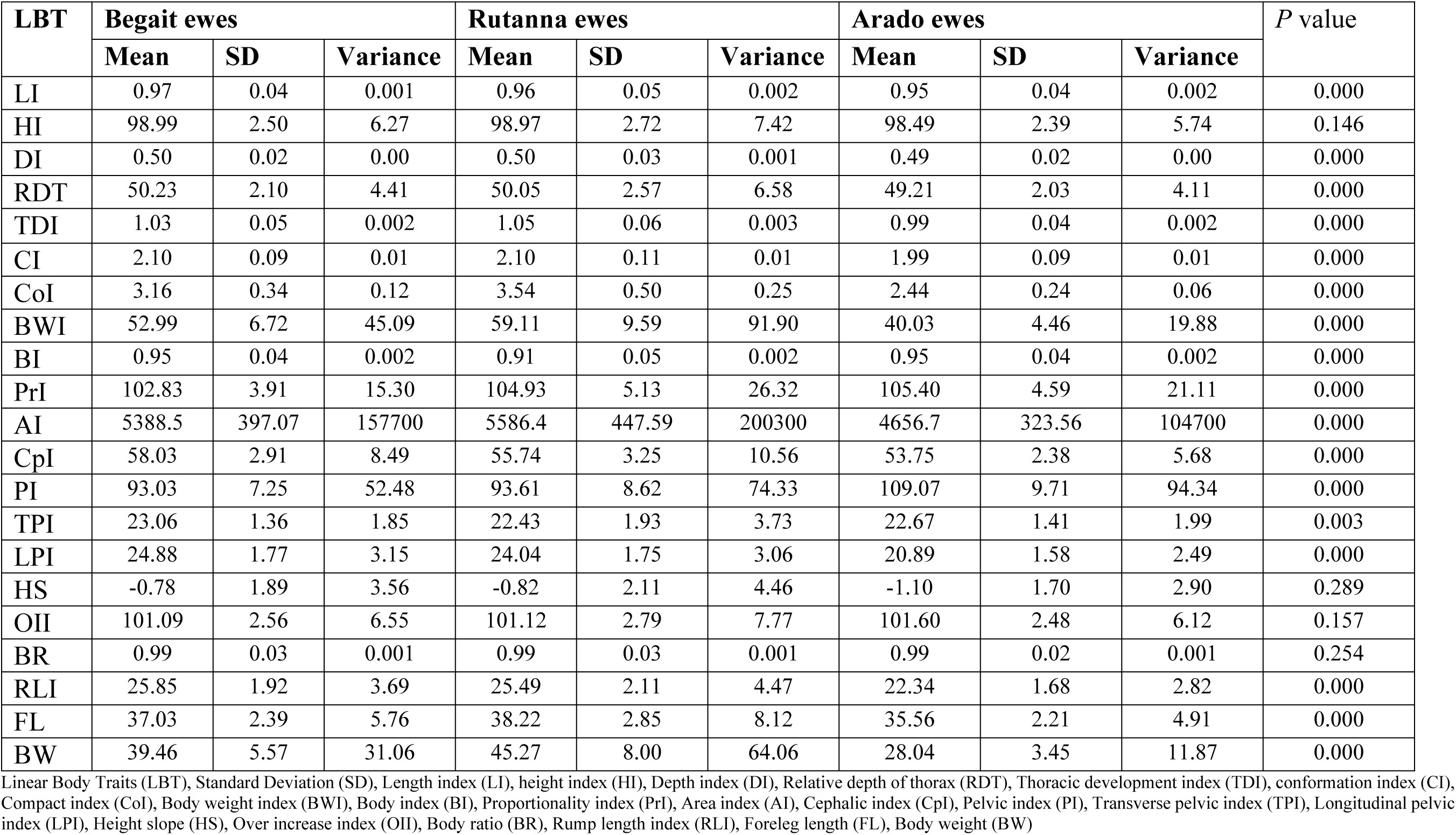
Biometric or morphometric or body structural indices of indigenous ewe populations.

| LBT | Begait ewes |  |  | Rutanna ewes |  |  | Arado ewes |  |  | P value |
| --- | --- | --- | --- | --- | --- | --- | --- | --- | --- | --- |
|  | Mean | SD | Variance | Mean | SD | Variance | Mean | SD | Variance |  |
| LI | 0.97 | 0.04 | 0.001 | 0.96 | 0.05 | 0.002 | 0.95 | 0.04 | 0.002 | 0.000 |
| HI | 98.99 | 2.50 | 6.27 | 98.97 | 2.72 | 7.42 | 98.49 | 2.39 | 5.74 | 0.146 |
| DI | 0.50 | 0.02 | 0.00 | 0.50 | 0.03 | 0.001 | 0.49 | 0.02 | 0.00 | 0.000 |
| RDT | 50.23 | 2.10 | 4.41 | 50.05 | 2.57 | 6.58 | 49.21 | 2.03 | 4.11 | 0.000 |
| TDI | 1.03 | 0.05 | 0.002 | 1.05 | 0.06 | 0.003 | 0.99 | 0.04 | 0.002 | 0.000 |
| CI | 2.10 | 0.09 | 0.01 | 2.10 | 0.11 | 0.01 | 1.99 | 0.09 | 0.01 | 0.000 |
| CoI | 3.16 | 0.34 | 0.12 | 3.54 | 0.50 | 0.25 | 2.44 | 0.24 | 0.06 | 0.000 |
| BWI | 52.99 | 6.72 | 45.09 | 59.11 | 9.59 | 91.90 | 40.03 | 4.46 | 19.88 | 0.000 |
| BI | 0.95 | 0.04 | 0.002 | 0.91 | 0.05 | 0.002 | 0.95 | 0.04 | 0.002 | 0.000 |
| PrI | 102.83 | 3.91 | 15.30 | 104.93 | 5.13 | 26.32 | 105.40 | 4.59 | 21.11 | 0.000 |
| AI | 5388.5 | 397.07 | 157700 | 5586.4 | 447.59 | 200300 | 4656.7 | 323.56 | 104700 | 0.000 |
| CpI | 58.03 | 2.91 | 8.49 | 55.74 | 3.25 | 10.56 | 53.75 | 2.38 | 5.68 | 0.000 |
| PI | 93.03 | 7.25 | 52.48 | 93.61 | 8.62 | 74.33 | 109.07 | 9.71 | 94.34 | 0.000 |
| TPI | 23.06 | 1.36 | 1.85 | 22.43 | 1.93 | 3.73 | 22.67 | 1.41 | 1.99 | 0.003 |
| LPI | 24.88 | 1.77 | 3.15 | 24.04 | 1.75 | 3.06 | 20.89 | 1.58 | 2.49 | 0.000 |
| HS | -0.78 | 1.89 | 3.56 | -0.82 | 2.11 | 4.46 | -1.10 | 1.70 | 2.90 | 0.289 |
| OII | 101.09 | 2.56 | 6.55 | 101.12 | 2.79 | 7.77 | 101.60 | 2.48 | 6.12 | 0.157 |
| BR | 0.99 | 0.03 | 0.001 | 0.99 | 0.03 | 0.001 | 0.99 | 0.02 | 0.001 | 0.254 |
| RLI | 25.85 | 1.92 | 3.69 | 25.49 | 2.11 | 4.47 | 22.34 | 1.68 | 2.82 | 0.000 |
| FL | 37.03 | 2.39 | 5.76 | 38.22 | 2.85 | 8.12 | 35.56 | 2.21 | 4.91 | 0.000 |
| BW | 39.46 | 5.57 | 31.06 | 45.27 | 8.00 | 64.06 | 28.04 | 3.45 | 11.87 | 0.000 |

The 49.79% of the height at withers (HW) of Begait ewes, 49.98% of the HW of Rutanna ewes and 50.81% of the HW of Arado ewes were their foreleg length proportions. These proportions of the foreleg lengths are much lower than the foreleg length proportions of the WAD (63.03%) and Yankasa (62.67%) sheep (Salako, 2006) due to the long depths of chest depths exhibited in Begait, Rutanna and Arado ewes which could be influenced by genotype, environment, the interaction effect of genotype and environment, sample size and field measurement errors. Rutanna ewes (3.54) exhibited much higher compact index (CoI) followed by Begait (3.16) ewes (Table, 9). Hence, both Begait and Rutanna ewe populations are meat type animals whilst Arado ewes (2.44) are more suitable for milk production. However, Begait ewes could be used as dual purpose animals because its CoI is close to the minimum standard (>3.15). The body weight index (BWI) of the Rutanna ewes (59.11) was higher than the BWI of the Begait (52.99) and Arado (40.03) ewes (Table 9) due to mainly their genetic variations. The BWI gives a reliable evaluation of animals to animal buyers because the measurement traits are associated with the production characteristics (Chacon *et al*., 2011). The longitudinal pelvic index (LPI) of Arado ewes (20.89) was also lower than the LPI of the Begait (24.88) and Rutanna (24.04) ewes (Table 9) which revealed that Begait and Rutanna ewes are classified as meat type animals.

The height slope (HS) of the three ewe populations revealed that the RuH of the ewe populations was longer than their HW. The HS of Begait, Rutanna and Arado ewes are not similar with the HS of Barbados Black Belly Cross (BC), Garut Local (GL), Garut Composite (GC), Sumatra Composite (SC) and St. Croix Cross (SCC) Sheep in Indonesia (Handiwirawan *et al*., 2011) and HS of WAD (2.54) and Yankasa (3.59) sheep in Nigeria (Salako, 2006) because the HS of Begait, Rutanna and Arado ewes were negative (HW<RuH). The differences in HS could be exhibited due to genotype, environment, interaction effect of genotype-environment, sample size and errors in measurements. The LI of Begait, Rutanna and Arado ewes are similar with the LI of GL (0.99) and SC (0.95) Sheep. Moreover, DI of Begait, Rutanna and Arado ewes are similar with the DI of BC (0.52) and GL (0.49) Sheep, and FL of Begait, Rutanna and Arado ewes are similar with the FL of SC (36.49) Sheep (Handiwirawan *et al*., 2011). The DI of Begait, Rutanna and Arado ewes are in agreement with the DI of WAD (0.53) and Yankasa (0.52) sheep (Salako, 2006), and the LI and FL of Begait, Rutanna and Arado ewes are also similar with the LI (0.93) and FL (36.16) of the Yankasa sheep (Salako, 2006).

According to Latorre *et al*. (2011), Body Index (BI) explains the relative capacity of the animal format. The present BI of Begait, Rutanna and Arado ewes are not similar with the BI of the Corriedale (0.72) and Australian Merino (0.74) sheep (Latorre *et al*., 2011) due to mainly their genetic makeup. Moreover, the relative depth of thorax (RDT) of Begait and Rutanna ewes are not similar with the RDT of the Corriedale (45.6) and Australian Merino (49.9) but the RDT of Arado ewes is in line with the RDT of the Australian Merino sheep (Latorre *et al*., 2011). The RDT of Begait, Rutanna and Arado ewes was lower than the RDT of Batur (54.62) sheep (Ibrahim *et al*., 2023) due mainly to their genetic makeup.

Arado ewes exhibited poor (0.99) thoracic development index (TDI) compared to TDI of Begait ewes (1.03) and Rutanna ewes (1.05). The Arado ewes were also weaker animals because the conformation index (CI) was lower (1.99) whilst Begait and Rutanna ewes were traction type animals (meat type) because the CIs of both ewe populations was 2.10 (Table 9). The Begait and Rutanna ewes could travel long distances in search of feed and water. According to the compact index (CoI) values, Rutanna and Begait ewes are categorized as mutton animals, however, Begait ewes could also serve as dual purpose animals. Three of the ewe populations were *longiline* because the body indices were higher than the BI standard set (BI>0.90). The rump length index (RLI) of the Arado ewes (22.34) was lower than the RLI of the Begait (25.85) and Rutanna (25.49) ewes, however, the pelvic index (PI) of the Arado ewes (109.07) was higher than the PI of the Begait (93.03) and Rutanna (93.61) ewes (Table 9). The higher PI value of the Arado ewes indicates that the breed has a good reproductive fitness because the PI is closely related to reproductive fitness (Israel *et al*., 2013).

The proportionality index (PrI) of the Wonosobo sheep (91.68) and Batur sheep (88.29) of Indonesia (Ibrahim *et al*., 2023) are much lower than the mean PrI of Begait, Rutanna and Arado ewes. Although the thoracic development index (TDI) of Arado ewes is in line with the TDI (0.99) of the Koroji sheep (Dauda, 2018), the present mean LI, PI, PrI and AI of Begait, Rutanna and Arado ewes are not similar with the LI (1.63), PI (81.89), PrI (160.18) and AI (4552.80) of the Koroji sheep in Nigeria (Dauda, 2018a). Moreover, the mean TDI of the Begait, Rutanna and Arado ewes are similar with the Balami sheep in Nigeria (Dauda, 2018b), however, the means of LI, PI, PrI, and AI of the Begait, Rutanna and Arado ewes are not similar with the LI (1.65), PI (73.92), PrI (155.22), and AI (653.80) of the Balami sheep in Nigeria (Dauda, 2018b). The differences in most of the indices of the Begait, Rutanna and Arado ewes and the indices of the two Nigerian sheep populations (Koroji and Balami) could be due to genotype, environment, interaction effect of genotype and environment and their sample size used.

There were little or no correlations (*P*>0.05) in the morphometric indices of Begait ewes which comprised of LI*CoI (0.117), HI*CoI (-0.014), HI*BWI (-0.098), HI*BI (0.058), LI*AI (0.055), DI*AI (0.023), RDT*AI (0.025), TDI*AI (-0.118), CI*AI (-0.123), BI*AI (0.145), PrI*AI (-0.046), CoI*BR (-0.009), BWI*BR (-0.091), BI*BR (0.036), CpI*BR (-0.137), and PI*BR (0.017) whilst the morphometric indices of Rutanna ewes with little or no correlations (*P*>0.05) comprised of CpI*LI (0.066), PI*LI (0.118), LPI*LI (0.115), HS*LI (-0.173), CoI*HI (0.005), BWI*HI (-0.074), BI*HI (0.027), CpI*HI (0.085), PI*HI (0.078), TPI*HI (0.137), LPI*HI (0.063), RLI*HI (-0.169), BW*HI (0.013), AI*CI (0.135), CpI*CI (-0.085), PI*CI (0.053), TPI*CI (0.081), LPI*CI (0.016), CpI*CoI (-0.082), PI*CoI (0.105), TPI*CoI (0.038), LPI*CoI (-0.076), HS*CoI (0.004), OII*CoI (-0.013), BR*CoI (-0.006), FL*CoI (0.053), CpI*BWI (-0.053), PI*BWI (0.152), TPI*BWI (0.148), LPI*BWI (-0.009), HS*BWI (-0.073), OII*BWI (0.066), BR*BWI (-0.085), AI*BI (0.041), CpI*BI (0.137), PI*BI (0.040), TPI*BI (0.115), LPI*BI (0.089), HS*BI (0.034), OII*BI (-0.027), BR*BI (0.029), FL*BI (0.037), RLI*BR (-0.166), and BW*BR (0.004) (Table 10). Moreover, the morphometric indices of Arado ewes with little or no correlations (*P*>0.05) comprised of AI*LI (0.114), CpI*LI (0.011), PI*LI (0.118), TDI*HI (-0.080), CI*HI (-0.092), CoI*HI (0.040), BWI*HI (-0.048), CpI*HI (-0.034), PI*HI (-0.020), RLI*HI (0.081), BW*HI (0.052), AI*CI (0.065), CpI*CI (-0.116), PI*CI (0.098), HS*CI (-0.092), OII*CI (0.093), BR*CI (-0.100), RLI*CI (-0.056), CpI*CoI (-0.045), LPI*CoI (-0.022), HS*CoI (0.029), OII*CoI (-0.043), BR*CoI (0.026), RLT*CoI (-0.155), FL*CoI (0.107), CpI*BWI (-0.053), LPI*BWI (0.022), HS*BWI (-0.056), OII*BWI (0.045), BR*BWI (-0.058), AI*BI (0.053), CpI*BI (0.130), PI*BI (0.029), TPI*BI (0.011), LPI*BI (-0.035), HI*BI (-0.151), BR*BI (-0.139), FL*BI (-0.084), RLI*BR (0.082), and BW*BR (0.037) (Table 11).

**Table 10.** Pearson’s correlation coefficient of the morphometric indices and body weights of Begait and Rutanna ewe populations (Top matrix- Begait and Bottom matrix - Rutanna)

| Indices | LI | HI | DI | RD T | TDI | CI | CoI | BWI | BI | PrI | AI | CpI | PI | TPI | LPI | HS | OII | BR | RLI | FL | BW |
| --- | --- | --- | --- | --- | --- | --- | --- | --- | --- | --- | --- | --- | --- | --- | --- | --- | --- | --- | --- | --- | --- |
| LI | 1 | -0.222** | 0.546** | 0.545** | 0.423** | 0.437** | 0.117 | 0.282** | 0.426** | -0.997** | 0.055 | -0.138 | 0.083 | 0.207** | 0.082 | -0.220** | 0.221** | -0.226** | -0.361** | -0.611** | 0.126 |
| HI | -0.189* | 1 | -0.270** | -0.272** | -0.251** | -0.257** | -0.014 | -0.098 | 0.058 | 0.225** | 0.218** | -0.158* | 0.021 | 0.303** | 0.212** | 0.999** | -0.999** | 0.994** | -0.020 | 0.368** | 0.000 |
| DI | 0.451** | -0.258** | 1 | 0.992** | 0.762** | 0.761** | 0.256** | 0.544** | -0.291** | -0.550** | 0.023 | -0.068 | 0.074 | 0.252** | 0.126 | -0.267** | 0.274** | -0.260** | -0.071 | -0.784** | 0.414** |
| RD T | 0.470** | -0.253** | 0.995** | 1 | 0.768** | 0.767** | 0.261** | 0.551** | -0.298** | -0.550** | 0.025 | -0.065 | 0.080 | 0.255** | 0.123 | -0.268** | 0.274** | -0.260** | -0.075 | -0.788** | 0.420** |
| TDI | 0.472** | -0.199* | 0.623** | 0.629** | 1 | 0.997** | 0.471** | 0.659** | -0.633** | -0.432** | -0.118 | 0.013 | 0.090 | 0.323** | 0.168* | -0.247** | 0.252** | -0.234** | 0.023 | -0.687** | 0.495** |
| CI | 0.471** | -0.202* | 0.628** | 0.634** | 0.998** | 1 | 0.474** | 0.662** | -0.622** | -0.445** | -0.123 | 0.008 | 0.083 | 0.322** | 0.174* | -0.253** | 0.258** | -0.240** | 0.024 | -0.694** | 0.494** |
| CoI | 0.284** | 0.005 | 0.222* | 0.235** | 0.637** | 0.648** | 1 | 0.949** | -0.380** | -0.121 | 0.461** | -0.033 | 0.195* | 0.176* | -0.070 | -0.022 | 0.013 | -0.009 | -0.129 | 0.062 | 0.951** |
| BWI | 0.401** | -0.074 | 0.515** | 0.527** | 0.757** | 0.767** | 0.949** | 1 | -0.424** | -0.287** | 0.400** | -0.054 | 0.190* | 0.232** | -0.018 | -0.105 | 0.098 | -0.091 | -0.136 | -0.207** | 0.957** |
| BI | 0.438** | 0.027 | -0.241** | -0.231** | -0.579** | -0.581** | -0.389** | -0.409** | 1 | -0.419** | 0.145 | -0.127 | -0.018 | -0.145 | -0.099 | 0.055 | -0.059 | 0.036 | -0.330** | 0.151 | -0.398** |
| PrI | -0.997** | 0.186* | -0.443** | -0.462** | -0.470** | -0.469** | -0.291** | -0.404** | -0.441** | 1 | -0.046 | 0.141 | -0.084 | -0.215** | -0.087 | 0.223** | -0.224** | 0.229** | 0.357** | 0.621** | -0.128 |
| AI | 0.189* | 0.254** | -0.005 | 0.013 | 0.127 | 0.135 | 0.576** | 0.505** | 0.041 | -0.198* | 1 | -0.064 | 0.055 | -0.065 | -0.116 | 0.209** | -0.221** | 0.221** | -0.212** | 0.544** | 0.616** |
| CpI | 0.066 | 0.085 | 0.046 | 0.049 | -0.085 | -0.085 | -0.082 | -0.053 | 0.137 | -0.072 | -0.019 | 1 | -0.151 | -0.196* | 0.008 | -0.152 | 0.155 | -0.137 | 0.135 | 0.043 | -0.043 |
| PI | 0.118 | 0.078 | 0.198* | 0.190* | 0.049 | 0.053 | 0.105 | 0.152 | 0.040 | -0.122 | 0.142 | -0.071 | 1 | 0.497** | -0.698** | 0.016 | -0.016 | 0.017 | -0.722** | -0.041 | 0.177* |
| TPI | 0.232** | 0.137 | 0.344** | 0.350** | 0.086 | 0.081 | 0.038 | 0.148 | 0.115 | -0.238** | 0.028 | 0.210* | 0.646** | 1 | 0.269** | 0.301** | -0.301** | 0.302** | 0.042 | -0.260** | 0.161* |
| LPI | 0.11<br>5 | 0.063 | 0.15<br>4 | 0.16<br>9 | 0.026 | 0.016 | -<br>0.076 | -<br>0.009 | 0.089 | -<br>0.117 | -<br>0.12<br>8 | 0.33<br>5** | -<br>0.45<br>8** | 0.375<br>** | 1 | 0.218<br>** | -<br>0.217<br>** | 0.216<br>** | 0.842<br>** | -<br>0.171<br>* | -<br>0.063 |
| HS | -<br>0.17<br>3 | 0.999<br>** | -<br>0.25<br>2** | -<br>0.24<br>6** | -<br>0.191<br>* | -<br>0.194<br>* | 0.004 | -<br>0.073 | 0.034 | 0.170 | 0.24<br>3** | 0.08<br>1 | 0.08<br>0 | 0.136 | 0.06<br>0 | 1 | -<br>0.999<br>** | 0.993<br>** | -<br>0.015 | 0.360<br>** | -<br>0.009 |
| OII | 0.18<br>6* | -<br>0.999<br>** | 0.25<br>5** | 0.25<br>0** | 0.196<br>* | 0.199<br>* | -<br>0.013 | 0.066 | -<br>0.027 | -<br>0.183<br>* | -<br>0.25<br>8** | -<br>0.08<br>9 | -<br>0.07<br>9 | -<br>0.139 | 0.06<br>5 | -<br>0.999<br>** | 1 | -<br>0.993<br>** | 0.016 | -<br>0.371<br>** | 0.000 |
| BR | -<br>0.20<br>7* | 0.995<br>** | -<br>0.26<br>0** | -<br>0.25<br>7** | -<br>0.217<br>* | -<br>0.220<br>* | -<br>0.006 | -<br>0.085 | 0.029 | 0.206<br>* | 0.24<br>8** | 0.08<br>2 | 0.09<br>1 | 0.138 | 0.05<br>1 | 0.994<br>** | -<br>0.994<br>** | 1 | -<br>0.012 | 0.364<br>** | 0.008 |
| RLI | -<br>0.42<br>5** | -0.169 | -<br>0.04<br>4 | -<br>0.04<br>4 | -<br>0.194<br>* | -<br>0.201<br>* | -<br>0.241<br>** | -<br>0.224<br>* | -<br>0.187<br>* | 0.426<br>** | -<br>0.31<br>5** | 0.22<br>3* | -<br>0.50<br>5** | 0.137 | 0.78<br>6** | -<br>0.181<br>* | 0.170 | -<br>0.166 | 1 | 0.030 | -<br>0.131 |
| FL | -<br>0.54<br>6** | 0.369<br>** | -<br>0.83<br>0** | -<br>0.83<br>0** | -<br>0.518<br>** | -<br>0.517<br>** | 0.053 | -<br>0.223<br>* | 0.037 | 0.537<br>** | 0.46<br>9** | -<br>0.06<br>7 | -<br>0.09<br>1 | -<br>0.302<br>** | -<br>0.22<br>6* | 0.353<br>** | -<br>0.368<br>** | 0.375<br>** | -<br>0.003 | 1 | 0.046 |
| BW | 0.27<br>7** | 0.013 | 0.40<br>7** | 0.41<br>9** | 0.655<br>** | 0.667<br>** | 0.958<br>** | 0.972<br>** | -<br>0.419<br>** | -<br>0.282<br>** | 0.66<br>2** | -<br>0.06<br>4 | 0.15<br>7 | 0.108 | -<br>0.05<br>8 | 0.009 | -<br>0.020 | 0.004 | -<br>0.223<br>* | -<br>0.024 | 1 |

\*\* . Correlation is significant at the 0.01 level (2-tailed); \* . Correlation is significant at the 0.05 level (2-tailed)..., Length index (LI), height index (HI), Depth index (DI), Relative depth of thorax (RDT), Thoracic development index (TDI), conformation index (CI), Compact index (CoI), Body weight index (BWI), Body index (BI), Proportionality index (PrI), Area index (AI), Cephalic index (CpI), Pelvic index (PI), Transverse pelvic index (TPI), Longitudinal pelvic index (LPI), Height slope (HS), Over increase index (OII), Body ratio (BR), Rump length index (RLI), Foreleg length (FL), Body weight (BW)

**Table 11.**
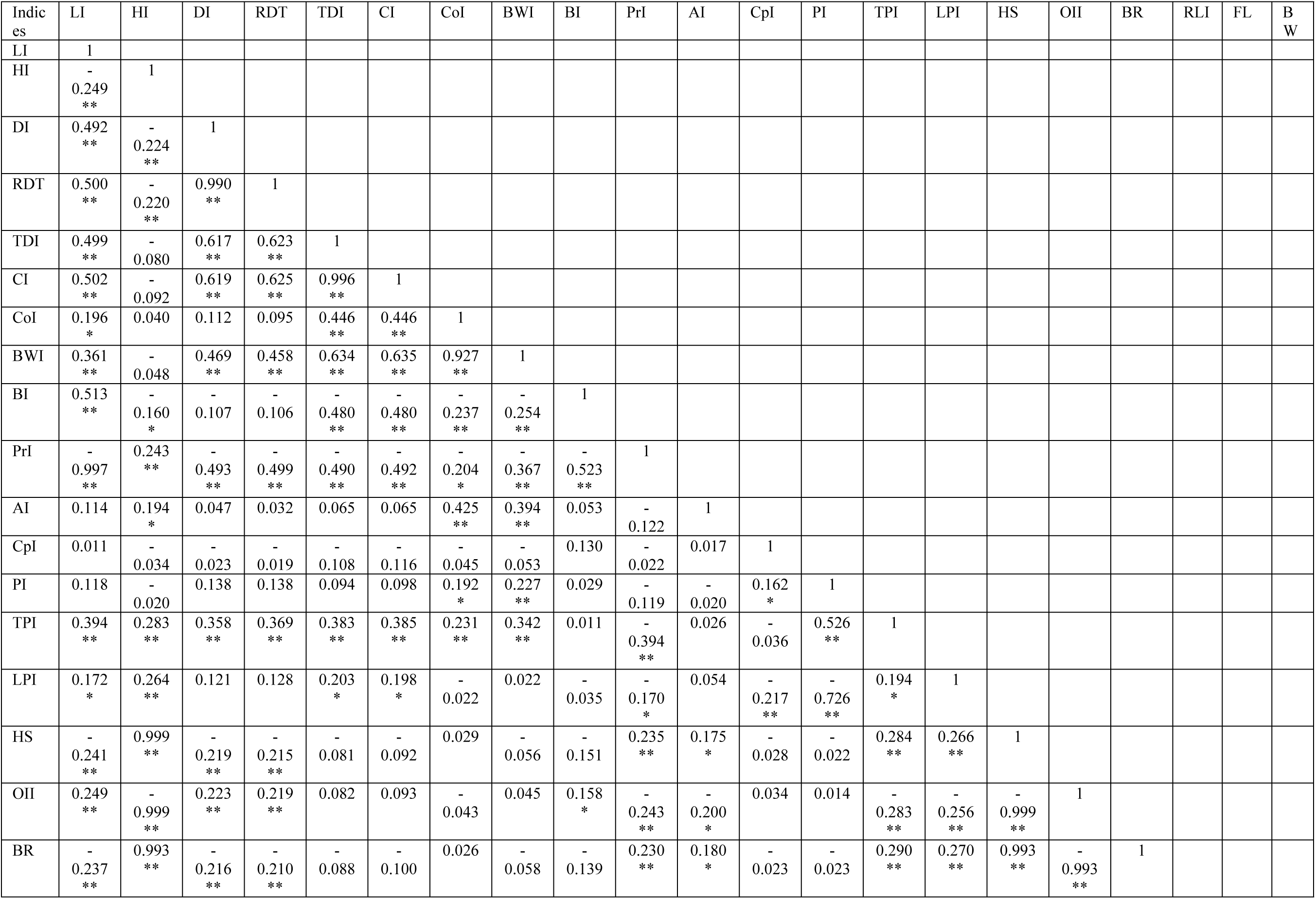
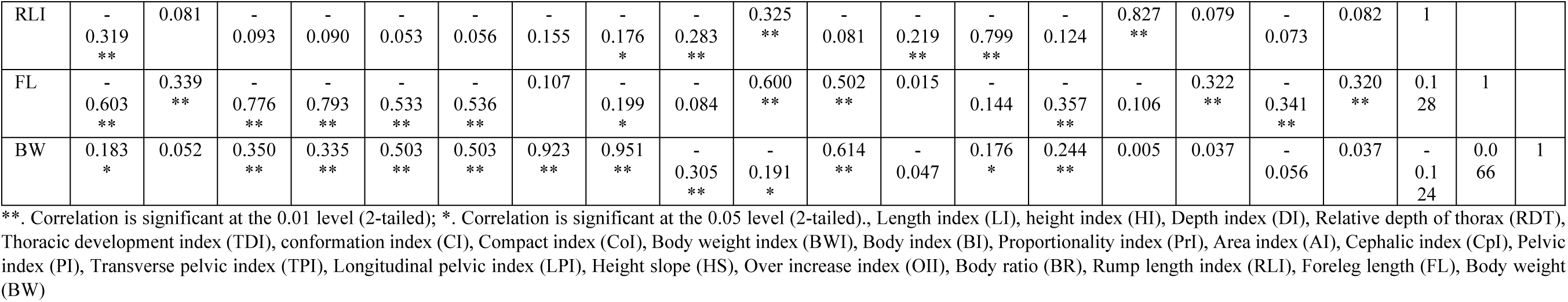
Pearson’s correlation coefficient of the morphometric indices and body weight of the indigenous Arado ewe population.

There were also little or no correlation among the morphometric indices and body weight of Begait ewes which comprised of LI*BW (0.126), HI*BW (0.000), PrI*BW (-0.128), CpI*BW (-0.043), LPI*BW (-0.063), HS*BW (-0.009), OII*BW (0.000), BR*BW (0.008), RLI*BW (-0.131), and FL*BW (0.046) whilst that of Rutanna ewes comprised of BW*HI (0.013), BW*CpI (-0.064), BW*PI (0.157), BW*TPI (0.108), BW*LPI (-0.058), BW*HS (0.009), BW*OII (-0.020), BW*BR (0.004), and BW*FL (-0.024) (Table 10). BW*HI (0.052), BW*CpI (-0.047), BW*LPI (0.005), BW*HS (0.037), BW*OII (-0.056), BW*BR (0.037), BW*RLI (-0.124), and BW*FL (0.066) were with little or no correlation among the morphometric indices and body weight of Arado ewes (Table 11).

Total variance explained (Table 12) and rotated component matrix (Table 13) of the different morphometric indices of each ewe populations are clearly presented. Morphometric indices with total initial eigenvalues greater than one were extracted as principal components (PCs). Accordingly, there were five PCs extracted which accounted for 86.7% of the total variance (Begait ewes), six PCs extracted which accounted for 91.5% of the total variance (Rutanna ewes) and five PCs extracted which accounted for 85.0% of the total variance (Arado ewes); the PCs were extracted from the different morphometric indices of the ewe populations (Tables 12 and 13). The sums of the rotated component matrix of the principal components (PCs) of the morphometric traits (Figure 2) and indices (Figure 3) of the ewe populations are also reported. The PC4 (3.401) of the morphometric indices of the Arado ewes revealed higher value of rotated component matrix compared to PC5 (0.809) of the Arado ewes, however, PC5 of the morphometric indices of the Begait and Rutanna ewes revealed a higher (2.908, 2.156) rotated component matrix as compared to their PC4 (0.641, 0.815), respectively (Figure 3).

**Figure 2.**
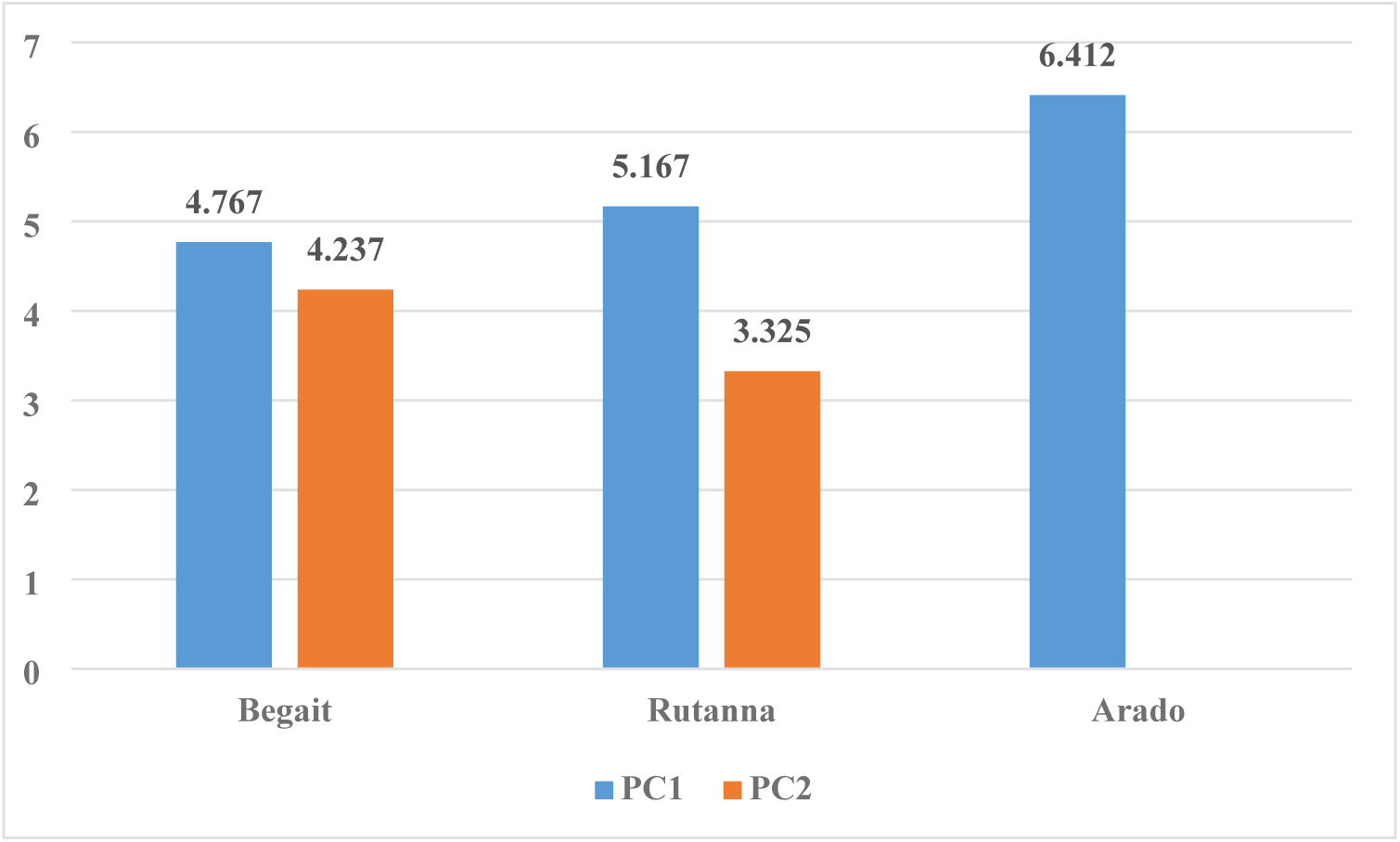
Sums of the Rotated Component Matrix of the Principal Components (PCs) of the Morphometric Traits of the Ewe Populations.

**Figure 3.**
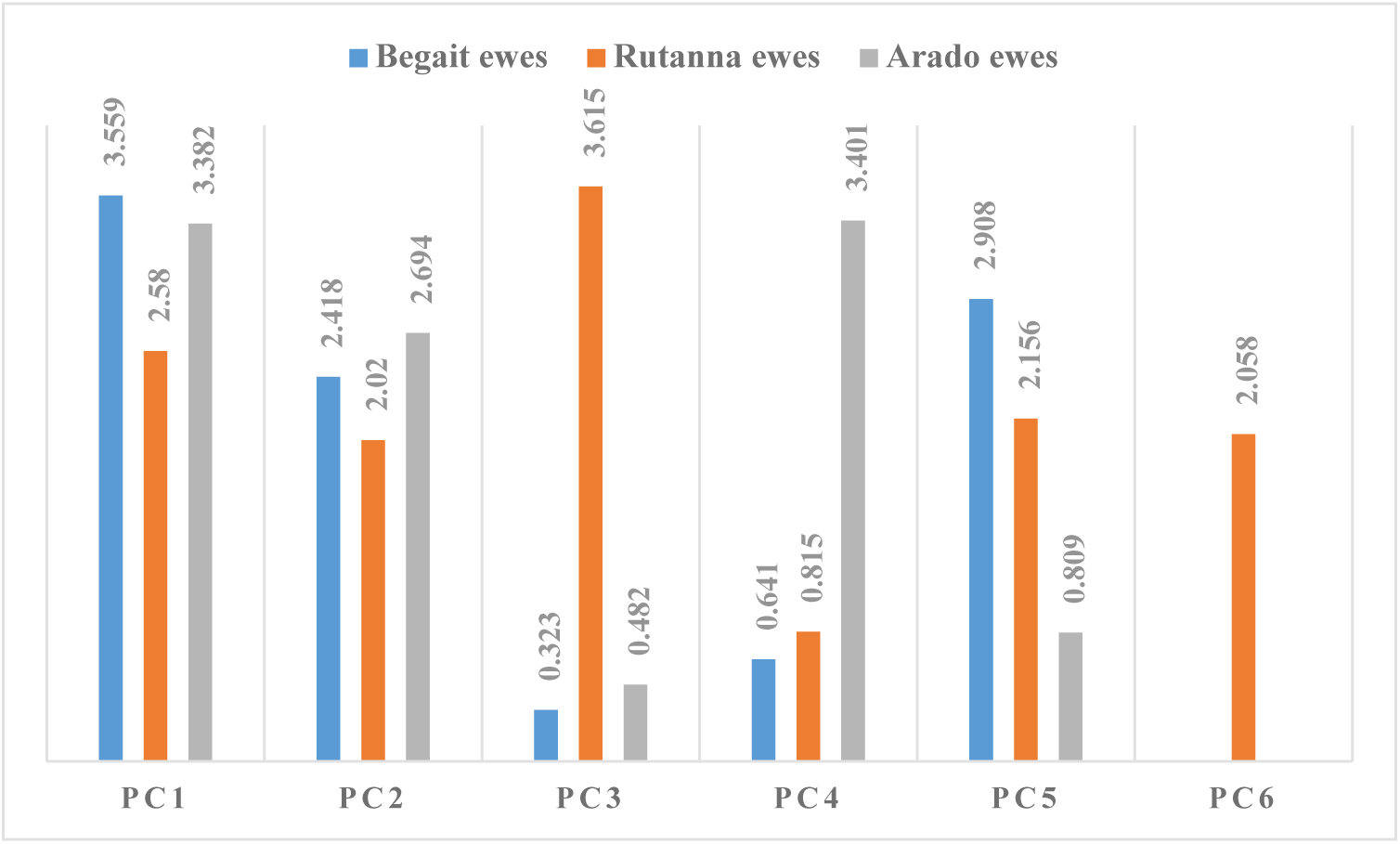
Sums of the Rotated Component Matrix of the Principal Components (PCs) of the Morphometric Indices of the Ewe Populations.

**Table 12.**
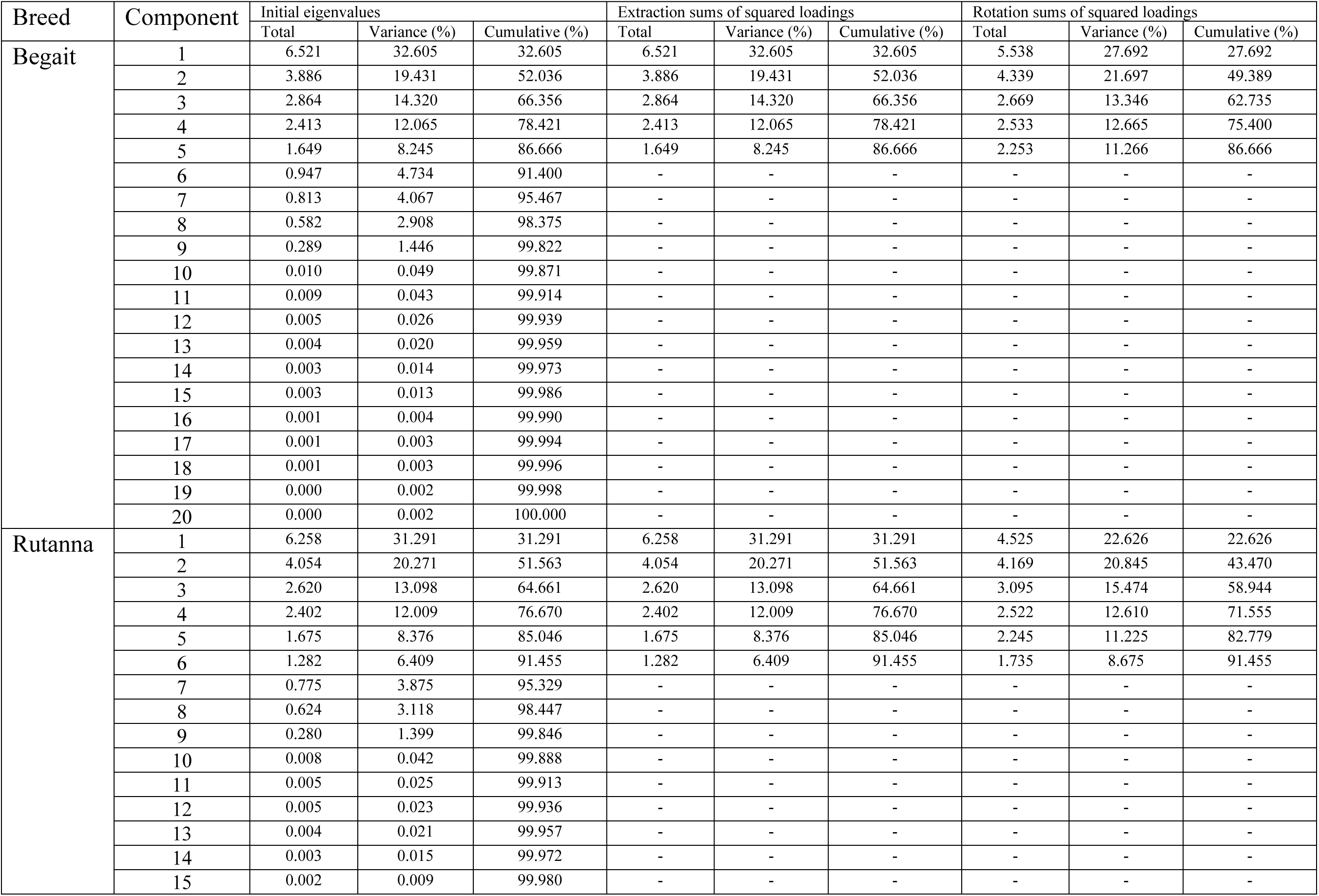
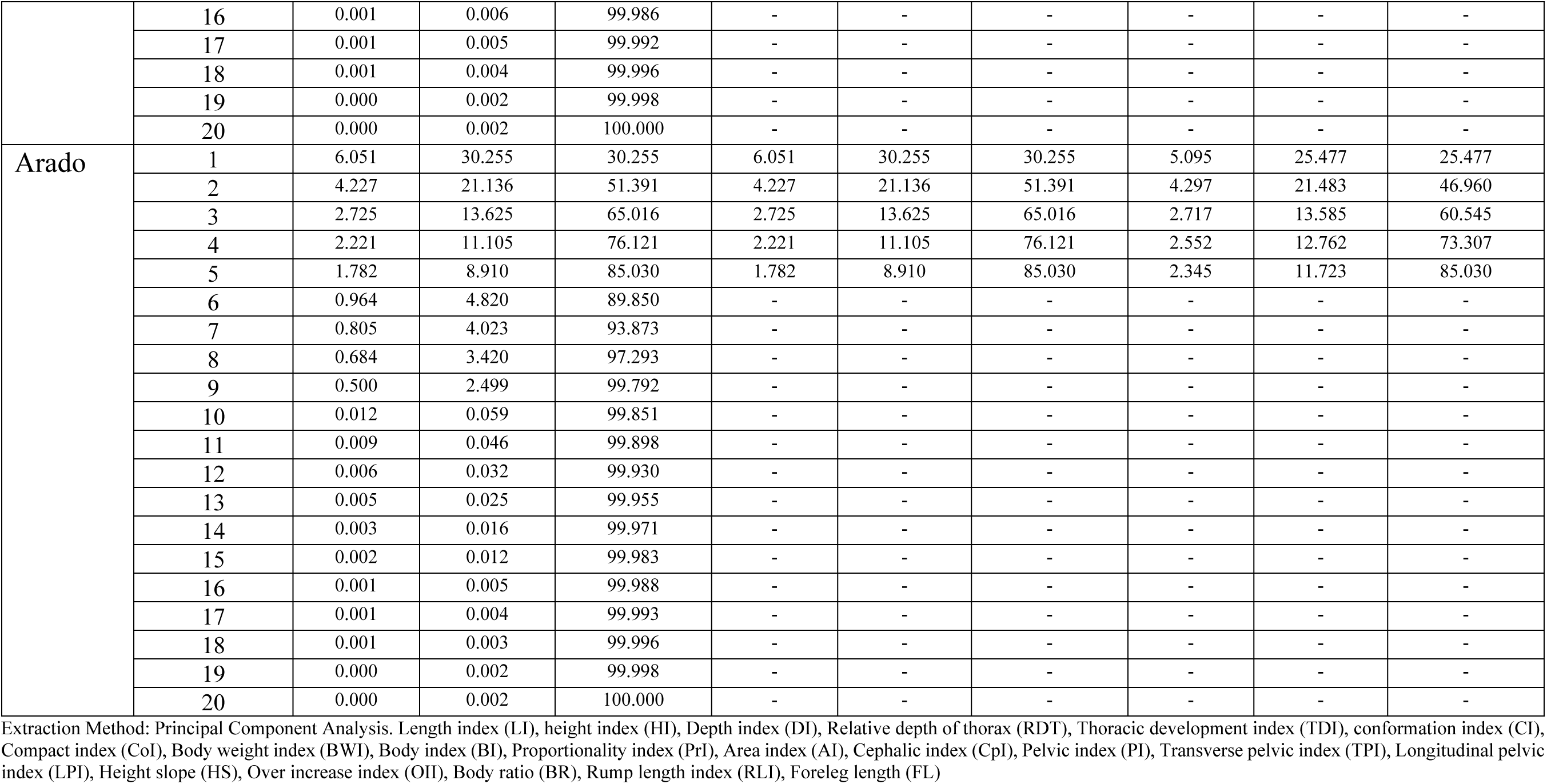
Total variance explained (biometric or morphometric indices)

**Table 13.** Rotated component matrix of different principal components (PCs) of morphometric indices of different sheep ewe populations.

| Breed | Traits | PCs |  |  |  |  |  |
| --- | --- | --- | --- | --- | --- | --- | --- |
|  |  | 1 | 2 | 3 | 4 | 5 | 6 |
| **Begait** | LI | 0.413 | -0.113 | 0.875 | -0.051 | 0.055 | - |
|  | HI | -0.169 | 0.969 | -0.050 | 0.043 | 0.063 | - |
|  | DI | 0.837 | -0.146 | 0.292 | 0.024 | 0.094 | - |
|  | RDT | 0.841 | -0.146 | 0.288 | 0.019 | 0.096 | - |
|  | TDI | 0.944 | -0.121 | -0.026 | 0.027 | 0.146 | - |
|  | CI | 0.944 | -0.126 | -0.013 | 0.033 | 0.146 | - |
|  | CoI | 0.393 | 0.012 | -0.143 | -0.124 | 0.824 | - |
|  | BWI | 0.615 | -0.035 | -0.027 | -0.099 | 0.741 | - |
|  | BI | -0.586 | 0.021 | 0.771 | -0.073 | -0.116 | - |
|  | PrI | -0.423 | 0.114 | -0.871 | 0.050 | -0.050 | - |
|  | AI | -0.218 | 0.162 | 0.174 | -0.044 | 0.843 | - |
|  | CpI | -0.060 | -0.255 | -0.243 | 0.160 | 0.013 | - |
|  | PI | 0.174 | 0.129 | -0.030 | -0.934 | -0.012 | - |
|  | TPI | 0.508 | 0.512 | 0.037 | -0.178 | -0.142 | - |
|  | LPI | 0.227 | 0.266 | 0.065 | 0.893 | -0.104 | - |
|  | HS | -0.164 | 0.969 | -0.050 | 0.049 | 0.053 | - |
|  | OII | 0.171 | -0.968 | 0.049 | -0.048 | -0.066 | - |
|  | BR | -0.155 | 0.965 | -0.064 | 0.050 | 0.068 | - |
|  | RLI | 0.055 | -0.015 | -0.370 | 0.869 | -0.150 | - |
|  | FL | -0.788 | 0.224 | -0.341 | -0.025 | 0.406 | - |
| **Rutanna** | LI | 0.416 | -0.109 | 0.219 | 0.864 | -0.004 | 0.023 |
|  | HI | -0.133 | 0.988 | 0.029 | -0.038 | 0.022 | 0.041 |
|  | DI | 0.872 | -0.154 | 0.086 | 0.105 | 0.083 | 0.223 |
|  | RDT | 0.871 | -0.149 | 0.101 | 0.123 | 0.096 | 0.218 |
|  | TDI | 0.783 | -0.101 | 0.520 | -0.007 | -0.108 | -0.129 |
|  | CI | 0.782 | -0.105 | 0.528 | -0.008 | -0.115 | -0.126 |
|  | CoI | 0.214 | 0.006 | 0.932 | -0.011 | -0.078 | -0.015 |
|  | BWI | 0.467 | -0.040 | 0.845 | 0.034 | -0.034 | 0.058 |
|  | BI | -0.426 | 0.001 | -0.325 | 0.806 | 0.110 | 0.131 |
|  | PrI | -0.407 | 0.108 | -0.229 | -0.865 | 0.000 | -0.029 |
|  | AI | -0.259 | 0.204 | 0.785 | 0.203 | -0.053 | 0.127 |
|  | CpI | -0.038 | 0.078 | -0.011 | 0.146 | 0.562 | 0.135 |
|  | PI | 0.084 | 0.062 | 0.067 | 0.045 | -0.385 | 0.903 |
|  | TPI | 0.245 | 0.131 | 0.021 | 0.102 | 0.396 | 0.822 |
|  | LPI | 0.166 | 0.072 | -0.047 | 0.058 | 0.940 | -0.134 |
|  | HS | -0.120 | 0.991 | 0.021 | -0.024 | 0.013 | 0.037 |
|  | OII | 0.132 | -0.988 | -0.036 | 0.037 | -0.025 | -0.043 |
|  | BR | -0.143 | 0.984 | 0.017 | -0.052 | 0.015 | 0.056 |
|  | RLI | -0.054 | -0.204 | -0.188 | -0.445 | 0.817 | -0.153 |
|  | FL | -0.872 | 0.245 | 0.280 | -0.258 | -0.096 | -0.087 |
| **Arado** | LI | 0.551 | -0.140 | -0.043 | 0.186 | 0.769 | - |
|  | HI | -0.108 | 0.981 | 0.062 | 0.047 | -0.084 | - |
|  | DI | 0.860 | -0.141 | 0.011 | 0.021 | 0.099 | - |
|  | RDT | 0.871 | -0.133 | 0.013 | 0.003 | 0.104 | - |
|  | TDI | 0.841 | -0.032 | 0.103 | 0.374 | -0.132 | - |
|  | CI | 0.842 | -0.043 | 0.100 | 0.375 | -0.132 | - |
|  | CoI | 0.188 | 0.019 | -0.086 | 0.896 | -0.088 | - |
|  | BWI | 0.494 | -0.035 | -0.076 | 0.803 | -0.044 | - |
|  | BI | -0.274 | -0.098 | -0.154 | -0.179 | 0.909 | - |
|  | PrI | -0.545 | 0.134 | 0.049 | -0.192 | -0.777 | - |
|  | AI | -0.190 | 0.159 | 0.048 | 0.748 | 0.205 | - |
|  | CpI | -0.092 | -0.017 | -0.327 | -0.056 | 0.108 | - |
|  | PI | 0.222 | 0.085 | -0.921 | 0.070 | -0.040 | - |
|  | TPI | 0.564 | 0.446 | -0.213 | 0.079 | 0.205 | - |
|  | LPI | 0.192 | 0.271 | 0.895 | -0.002 | 0.201 | - |
|  | HS | -0.100 | 0.982 | 0.061 | 0.030 | -0.076 | - |
|  | OII | 0.109 | -0.981 | -0.055 | -0.051 | 0.083 | - |
|  | BR | -0.102 | 0.981 | 0.062 | 0.027 | -0.065 | - |
|  | RLI | -0.088 | 0.031 | 0.914 | -0.130 | -0.216 | - |
|  | FL | -0.853 | 0.225 | 0.039 | 0.352 | -0.220 | - |

Extraction Method: Principal Component Analysis. Rotation Method: Varimax with Kaiser Normalization. Length index (LI), height index (HI), Depth index (DI), Relative depth of thorax (RDT), Thoracic development index (TDI), conformation index (CI), Compact index (CoI), Body weight index (BWI), Body index (BI), Proportionality index (PrI), Area index (AI), Cephalic index (CpI), Pelvic index (PI), Transverse pelvic index (TPI), Longitudinal pelvic index (LPI), Height slope (HS), Over increase index (OII), Body ratio (BR), Rump length index (RLI), Foreleg length (FL)

## CONCLUSION AND RECOMMENDATIONS

The principal component analysis (PCA) of the morphometric traits (09) extracted two principal components (PCs) which explained 63.91% of the total variance in the traits of Begait ewes, two PCs which explained 61.07% of the total variance in the traits of Rutanna ewes and one PC which explained 51.73% (not rotated variance) of the total variance in the traits of Arado ewes whilst the PCA of the morphometric indices (20) extracted five PCs which explained 86.66% of the total variance in the indices of Begait ewes, six PCs which explained 91.45% of the total variance in the indices of Rutanna ewes and five PCs which explained 85.03% of the total variance in the indices of Arado ewes.

Based on the biometric traits, Arado ewes are small in size whilst Rutanna and Begait ewes are larger in size and suitable for mutton production. The TDI, CI, CoI, BWI, AI and LPI largely confirmed that Rutanna and Begait ewes are important for mutton production.

The low-input extensive production system should be shifted to high-input intensive production system to boost their productivity. The fertility and growth rate of the Rutanna and Begait ewes should be strategically improved to boost their mutton yields. Genetic characterizations of the indigenous (Begait and Arado) and the transboundary (Rutanna) sheep populations should be complemented to confirm and distinguish the purpose of the populations.

## Author Contribution

Conceptualization: Teweldemedhn Mekonnen, Methodology: Teweldemedhn Mekonnen, Investigation: Teweldemedhn Mekonnen, Medhanye Araya and Solomon Tesfahun, Data curation: Teweldemedhn Mekonnen, Writing original draft preparation: Teweldemedhn Mekonnen, Writing-review and editing: Teweldemedhn Mekonnen and Medhanye Araya, Research administration: Teweldemedhn Mekonnen

## Data availability

The corresponding author will deliver the data upon request.

## Acknowledgement

The authors would greatly like to thank to Humera Agricultural Research Center of Tigray Agricultural Research Institute.

